# Forest edges and other semi-natural habitat edges increase wild bee species richness and habitat connectivity in intensively managed temperate landscapes

**DOI:** 10.1101/2024.07.05.602209

**Authors:** Markus A.K. Sydenham, Anders Nielsen, Yoko L. Dupont, Claus Rasmussen, Henning B. Madsen, Marianne S. Torvanger, Bastiaan Star

## Abstract

Pollinator conservation schemes are typically focused on conserving existing-, restoring degraded- or establishing new wild bee habitats. The effectiveness of such conservation schemes depends on the presence of dispersal corridors that allow habitat colonization by bees. Nonetheless, we lack an understanding of the role of semi-natural habitats edges on the connectivity of pollinator communities across intensively managed landscapes. Here, we use data from wild bee communities comprising 953 occurrences from 79 species of non-parasitic bees, sampled at 68 locations distributed across a Norwegian and a Danish landscape to show that the proportion of semi-natural habitat edges is positively correlated to bee species richness and habitat connectivity. Specifically, we found that wild bee species richness sampled along roadsides increased with the proportion of semi-natural habitat edges within1.5 km of the study sites and with local plant species richness. We combined maps showing the proportion of seminatural habitat edges with least cost path analysis to find the most likely dispersal route between our bee communities. We find that these least cost path lengths provide better models of bee species compositional similarity than geographic distance (|ΔAICc| > 2), suggesting that seminatural habitat edges act as dispersal corridors in intensively managed landscapes. However, we also find that compositional similarity between communities depend on site-specific plant species richness stressing the importance of improving the habitat quality of edge habitats if they are to function as dispersal corridors. We discuss potential management options for improving wild bee habitat conditions along seminatural habitat edges and illustrate how maps of least cost paths can be used to identify dispersal corridors between pollinator habitats of conservation priority. Maps of dispersal corridors can be used to direct wild bee habitat management actions along seminatural habitat edges to facilitate the dispersal of bees between larger grassland habitats.

## Introduction

Land use intensity and the associated loss of semi-natural habitats is a main driver of pollinator declines and threatens ecosystem functioning in the cultural landscape (Dicks et al., 2021). Pollinator diversity increases with habitat size (Steffan-Dewenter et al., 2006) and quality (Rollin et al., 2019) and conserving or restoring habitats is a central component of many pollinator conservation schemes (Senapathi et al., 2017). However, the species composition within habitats is dynamic (Leibold et al., 2004); species disperse across the landscape and colonize suitable patches of habitats (Franzén & Nilsson 2010) where they may remain or eventually drift to local extinction before the patch is potentially colonized again (Vellend, 2010; Hanski, 1998). Indeed, wild bee meta-population dynamics can be highly dynamic in terms of extinction and colonization events (Franzén & Nilsson 2010). The species diversity and functioning of pollinator communities within habitats therefore depends on habitat-specific environmental conditions that determine the number of individuals and species richness that a habitat can sustain (Krauss et al., 2009). Furthermore, spatial (Beduschi et al., 2018) and temporal (Griffin et al., 2017) connectedness to other habitat patches determine the species composition in habitats (Taylor et al., 1993). Mitigating insect declines and preserving ecological functions such as pollination therefore requires an understanding of how land use management affects the connectedness of pollinator communities and plant-pollinator interactions across landscapes (Cranmer et al., 2012). While it is well established that semi-natural habitat patches, often in the form of edges along land use types, provide important resources for pollinators in intensively managed landscapes (Eldegard et al., 2015; von Königslöw, 2021; Johansen et al., 2022) it is less well known how semi-natural edge habitats affect dispersal rates and thereby the connectivity of pollinator communities across intensively managed landscapes.

Pollinators can forage or nest in forest-field edges (Kells & Goulson 2003; Sydenham et al., 2014; Sõber et al., 2020), forest-shrubland edges (Glenny et al., 2023), grassland edges (Cole et al., 2015), road verges (Hopwood, 2008), and edges around sparsely vegetated areas such as quarries and other low productive areas (Heneberg & Bogusch 2020). Moreover, improving or introducing edge habitats in the form of flower strips (Haaland et al., 2011; von Königslöw et al., 2021) or hedgerows (Morandin & Kremen 2013) can provide pollinators with resources that are otherwise limiting in the landscape. By providing pollinators with nesting sites (Kells & Goulson 2003; Osborne et al., 2008; Morandin & Kremen 2013) and by guiding the flight direction of pollinators (Cranmer et al., 2012) the presence of semi-natural edge habitats may function as corridors and increase the dispersal rate of species through the landscape. Indeed, colonization of open habitats in a forested landscape by the solitary bee *Megachile rotundata* increases in the presence of open linear corridors between habitats (Griffin & Haddad 2021).

A lack of dispersal corridors among habitat patches can reduce species flow across the landscape, making local populations more vulnerable to stochastic fluctuations in population growth rates (Vellend 2010). Land use change can impact landscape connectivity if availability of semi-natural habitat patches is reduced along dispersal corridors of species. Hence, a reduction in semi-natural habitat in the wider landscape may result in dispersal barriers, increasing the rate of wild bee species turnover with geographic distance between habitat patches (Beduschi et al., 2018). For wild bees in power line clearings the likelihood of occurring within a site has also been shown to decrease with the distance to the nearest site where the species occurs (Sydenham et al., 2017). The similarity of bee communities can therefore be indicative of their connectedness. Least-cost-path analysis provides a useful framework for identifying the shortest route between two communities that minimizes the cost of travelling across the landscape and thereby reflects the most likely dispersal route between communities (Adriaensen et al., 2003). In least-cost-path analysis, the landscape is viewed as a resistance surface, where local environmental conditions can either increase or decrease the rate, or ‘cost’, of movement depending on the availability of habitats (Etherington, 2016). Least-cost-path analysis is frequently applied in landscape-genetics and in movement ecology to estimate effects from land use on the dispersal of species (Zeller et al., 2012). Least cost path analysis has also proven useful when studying patterns of species compositional similarity, where the accumulated habitat availability, or its inverse – the cost of movement, has been shown to be a better predictor of ant species turnover than geographic distance (Liu et al., 2018). Longer least cost path distances reduce the rate of patch colonization. As a result, colonization events cannot counter the effects of local extinctions and the species compositional similarity between communities drifts apart (Vellend, 2010). For pollinators, such as wild bees, the presence of semi-natural edge habitats could be expected to reduce the resistance of the landscape to movement because dispersing individuals can find nesting and foraging habitats within edge habitats (Kells & Goulson 2003). Edge habitats may therefore act as steppingstones along dispersal corridors between larger habitats (Menz et al., 2011) and thereby contribute to preserving metapopulation dynamics (Saura et al., 2014; Hanski 1998). However, to our knowledge, the potential contribution of habitat edges to wild bee habitat connectivity in intensively managed landscapes has not yet been formally investigated.

We use data from 68 wild bee communities sampled across a Norwegian and a Danish landscape to test whether semi-natural edge habitats increase the connectedness of wild bee communities and to illustrate how least cost path analyses can be used to identify important dispersal corridors for wild bees in intensively managed landscapes. We hypothesize that:

(1) Semi-natural habitat edges provide habitat resources for wild bees in managed landscapes. We test if wild bee species richness in semi-natural plant communities increases with the amount of semi-natural habitat edges within typical wild bee foraging ranges. Specifically, we test if wild bee species richness increased with the amount of (i) semi-natural habitat edges including forest edges, (ii) forest edges alone, (iii) or semi-natural habitat edges excluding forest edges. We define semi-natural edge habitats as edges around forest, grassland, shrubland, wetlands, and sparsely vegetated areas.
(2) Semi-natural habitat edges increase the connectivity of bee communities across the landscape. We test if the species compositional similarity and the number of shared species between pairs of sites decreases with the length and cost of least cost paths, where least cost paths are estimated from maps showing the proportion of seminatural habitat edges in the landscape. We compare the relationships between compositional similarity and least cost path lengths to that of geographic distances, the latter acts as a null model by assuming that species can disperse freely across the landscape. We estimate least cost paths between site-pairs using maps of the proportion of 10m pixels classified as semi-natural habitat edges (forest, grassland, shrubland, wetlands, and sparsely vegetated areas) or only forest edges, to define the resistance of pixels to movement.

## Methods

### Sampling

We sampled plant-bee interactions in 68 semi-natural, forb-dominated, plant communities (Fig. 1). We focused on wild bees because (1) they are considered central pollinators for many plant species (Willmer et al., 2017), (2) bees are central place foragers with restricted home ranges, and (3) even large bees often show limited movement between habitat patches (Franzén et al., 2010), making them ideal for studying impacts of land use on habitat connectivity. We used linear open landscape features such as roadsides as a model system and established one 50×2m transect for our surveys in each site. To cover the main flowering period, we sampled flower visiting bees once in May, June and July at each site. To standardize sampling times across sites and countries, timing of the first sampling was determined by the peak flowering of dandelions (*Taraxacum officinale*). All flower-visiting bees were collected from flowers and stored in 96% EtOH prior to identification. Species within the *Bombus sensu stricto* subgenus are cryptic and cannot be reliably identified manually. Specimens within the *B. sensu stricto* subgenus were treated as one morpho-species. Each transect observation lasted 30 minutes, adding 30 seconds per collected specimen, to account for handling time. To sample all target species when they were active, sampling only took place on days with temperatures > 15°C, local wind speed < 5 m/s, with little to no cloud cover and no rain. At each site we conducted a vegetation survey in July and recorded the occurrence of herbaceous plants in ten 1m^2^ vegetation survey plots placed regularly along the transect. As an indicator of local plant species richness we tallied the number of plant species within a site for which we had observed at least one interaction with wild bees across the 68 sites.

**Figure 1.**
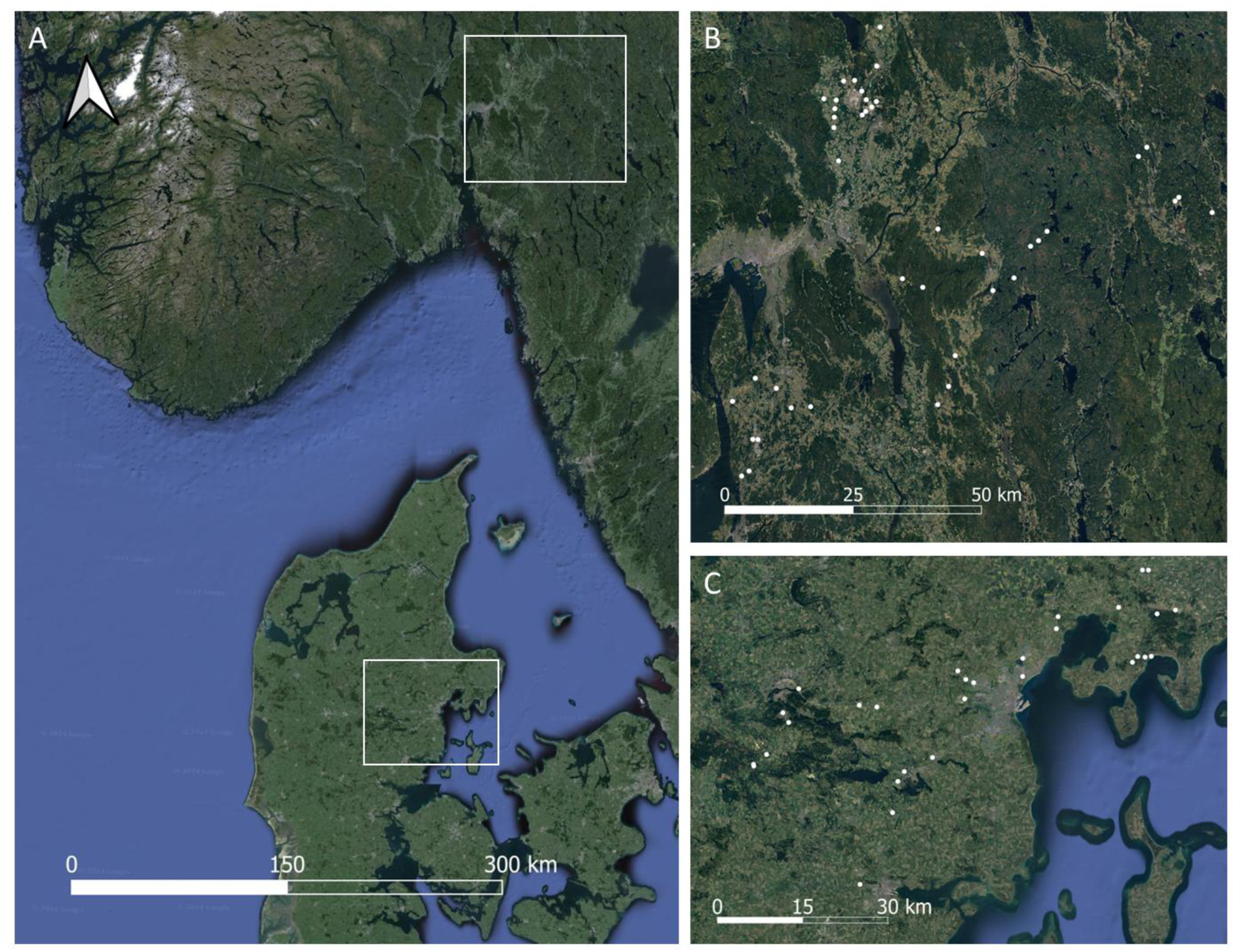
Wild bee communities were sampled at 68 roadsides and other linear habitats distributed along gradients of agricultural intensity within two landscapes (A) in southeastern Norway (B) and in western Denmark (C). Satellite imagery from Map data ©2024 Google via QGIS 2024

### Mapping edge habitats

We derived three habitat edge maps at a 10m resolution with maps showing (1) all semi-natural habitat edges, (2) forest edges only, and (3) all non-forest semi-natural edges. We used the CLC+ back bone land cover map from the European Environment Agency (© European Union, Copernicus Land Monitoring Service, 2022, European Environment Agency (EEA)). The CLC+ is a 10m resolution land cover map with pixels classified according into 11 land use classes: (1) Sealed; (2) Woody – needle leaved trees; (3) Woody – Broadleaved deciduous trees; (4) Woody – Broadleaved evergreen trees; (5) Low-growing woody plants (bushes, shrubs); (6) Permanent herbaceous; (7) Periodically herbaceous; (8) Lichens and mosses; (9) Non- and sparsely-vegetated; (10) Water; and (11) Snow and ice. We identified semi-natural edge habitats by using the as.lines, as.polygons, and rasterize functions from the terra package in R (Hijmans, 2023) to first polygonise all pixels classified as forest (CLC+ classes 2 and 3); low-growing woody plants (class 5); grassland (class 6); and non- and sparsely-vegetated (class (9), and then converting these polygons into lines which we rasterized to produce a raster of semi-natural habitat edges. We followed the same procedure to produce a raster map that included only forest edges (CLC+ classes 2 and 3), and one that included non-forest semi-natural habitat edges, i.e. by subtracting forest edges from the semi-natural habitat edge map prior to calculating the proportion of edge habitats.

### Statistical analyses

To test if semi-natural edges provide habitat for wild bees, we used a Poisson GLM and test if wild bee species richness increased with the proportion of semi-natural habitat edges, forest edges, or non-forest semi-natural edges within the surrounding landscape in R (R Core Team, 2023). We calculated the proportion of the area within circular buffers around each sampling location classified as habitat edges. To identify the spatial scale where the proportion of edge habitats had the strongest effect on bee species richness, we compared models where the proportion of edge habitats was calculated within buffers of 250m, 500m, 750m, 1000m, 1250, and 1500m and selected the model that best fitted the data, indicated by having the lowest AICc value. We ran models with the proportion of all semi-natural edge habitat, or as edge habitats defined by only including forest edges. For all models we included the species richness of flowering plants to control for site-specific differences in habitat quality. We used AICc to compare the best semi-natural edge habitat model against the forest edge habitat model and a model that only contained local plant species richness. We used DHARMa residuals (Hartig, 2022) to test for overdispersion and assess residual distributions from the models with the lowest AICc and likelihood ratio rest statistics to assess the statistical significance of edge habitat variables.

We calculated the wild bee species compositional similarity between study sites as 1-BC where BC was the Bray-Curtis dissimilarity calculated using the ‘vegdist’ function in Vegan (Oksanen et al., 2022). We excluded parasitic bees because their distributions are indirectly related to the environmental conditions that determine the distribution of their hosts. All BC dissimilarities were calculated on binary, presence-absence matrices. We assembled a data frame with one column defining the identity of the *i*th site, and another column defining the identity of the *j*th site. For each combination of the *i*th and *j*th sites we appended the wild bee species compositional similarity, in addition to the number of bee species shared between the two sites.

For each ith and jth site-pair we appended the geographic distance, and the least cost path lengths between sites using the maps of semi-natural edge habitats and forest edges. We calculated the geographic distance between all site pairs using the ‘distance’ function in terra (Hijmans, 2023) in R. To estimate least cost paths between site pairs, we used the ‘create_cs’ function with neighbors set to 8 in the leastcostpath (Lewis 2023) package to create a conductance surface from the 100m semi-natural habitat edge raster and from the 100m forest edge map, where we interpreted increasing proportions of semi-natural habitat edges and forest edges as increasing the permeability of the landscape to movement.

We adapted the function ‘create_lcp’ in leastcostpath (Lewis, 2023) to identify the least cost paths between all pairs of sites based on cost surface. We used the igraph package in R (Csárdi & Nepusz 2006; Csárdi et al., 2024) to identify least cost paths. We used the graph_from_adjency_matrix function with mode = ‘min’ and weighted = ‘TRUE’ to create a weighted graph from the conductance matrix between all cells in the cost surface. As in the ‘create_lcp’ function (Lewis, 2023) we 1/x transformed the weights to convert the edge weights from the conductance values into costs, with costs increasing exponentially as the proportion of edge habitats decreased. For each study site to all other study sites within the same country (Norway and Denmark) we then used the ‘shortest_paths’ function in igraph to identify all raster cells connecting site pairs along the shortest path and converted the coordinates of these cells into a spatial lines object (spatvector) and used the ‘perim’ function from terra (Hijmans, 2023) the calculate the length of the resulting lines (i.e. least cost paths). Least cost path lengths may not equate the cost of movement across the landscape (Etherington & Holland 2013). We therefore 1-x transformed the 100m habitat edge raster maps and extracted the resulting pixel values along each least cost path which we summed to obtain estimates of the total cost of movement along a path. Because we were interested in how land use conditions affect habitat connectivity we restricted our analyses to site-pair comparisons within the same country.

We used zero-inflated Beta GLMMs from the glmmTMB package (Mollie et al., 2017) to model the bee species compositional turnover with the site identity of the *i*_th_ and *j*_th_ sites as random intercept terms. Species compositional similarity showed a unomial response to the distance between sites which ranged from within one km up to 105 km (mean = 41.8 ± 24.4). To limit our analyses to the data where species compositional similarity decreased with distance, we conducted a sensitivity analysis to identify the geographic distance from which species compositional similarity increased with distance. We created a sequence of numbers from 10 to 105 with increments of 5 to create 20 subsets of our data where the uppermost distance between sites was restricted to within 5km increments from 10km to 105km. For each subset of the data, we used a zero-inflated Beta GLMM to model the species compositional similarity as a function of either the untransformed geographic distance between sites or as a function that included a second order polynomial transformation of geographic distance and calculated the difference in AICc values (Fig. S1. We then selected the largest distance (60km) where the polynomial and the untransformed models performed equally well (|ΔAICc| < 2), to include as much data as possible in subsequent tests. For site-pairs located less than 60km apart, the corresponding least cost paths when considering all semi-natural habitat edges (mean = 84km, min = 75km, max = 97km) and when considering only forest edges (mean = 85km, min = 76km, max = 104km) were considerably longer than the geographic distances. For all site-pairs within 60km of each other we used zero-inflated Beta GLMMs to model species compositional similarity and Generalized Poisson GLMMs to model the number of shared species. For all models, we included an interaction term between the plant species richness in the *i*th and *j*th to control for site-specific differences in plant diversity and its potentially confounding effect on species compositional similarity and the number of shared species. For both compositional similarity and the number of shared species, we used AICc to compare the goodness of fit of models with the least cost path length and path cost estimated from the proportion semi-natural habitat edges or from forest edges as fixed effects against models using geographic distance as fixed effect.

To illustrate how maps of least cost paths can aide in identifying important dispersal corridors between seminatural grasslands, we used georeferenced data on the location of 125 seminatural hay-meadows (EUNIS Annex I code: EUNIS R22) of high (“Klasse A”) or good (“Klasse B”) quality from within the extent of our Norwegian study sites. For the Danish case, we considered 314 georeferenced meadows within the spatial extent of our study sites, focusing on meadows that are included in the proposed Natura2000 action plan areas for 2022-2027 (The Danish Environmental Protection Agency 2023): 13 Xeric sand calcareous grasslands (EUNIS Annex I code: 6120);102 Semi-natural dry grasslands and scrubland facies on calcareous substrates (Festuco-Brometalia) [EUNIS Annex I code: 6210]; 212 Species-rich Nardus grasslands, on silicious substrates in mountain areas (and submountain areas in Continental Europe) [EUNIS Annex I code: 6230]; and 47 Molinia meadows on calcareous, peaty or clayey-silt-laden soils (Molinion caeruleae) [Annex I code: 6410). To focus on meadows that would be of likely high value for wild bees, we restricted the analyses to meadows of high (“I. Høj tilstand”) or good (“II. God tilstand”) condition and which were at least two hectares large, resulting in 23 EUNIS Annex I code 6210 grasslands and 54 EUNIS Annex I code 6230 grasslands. For the meadows within each region, we used the centroid location of each meadow to estimate least cost paths between all pairs of meadows based on proportion of seminatural habitat edges within 100m raster pixels.

## Results

We sampled a total of 953 occurrences of 79 species of non-parasitic wild bees distributed across 68 wild bee communities located in two regions, one in Denmark and one in Norway (Fig. 1). The sampled bees belonged to 21 genera within six the families: Melittidae; Andrenidae; Colletidae; Halictidae; Megachilidae; and Apidae. Bee species richness within sites ranged from two to 18 (mean = 9.34, standard deviation = 4.05). The species compositional similarity, or the proportion of species co-occurring in two sites, ranged from zero to 0.89 (mean = 0.35, s.d. = 0.15), while the number of shared species ranged from zero to 11 (mean = 3.49, s.d. = 1.95), when considering all site-pairs within each country. Values were similar when considering the 873 site-pairs located within 60km of each other: species compositional similarity ranging from zero to 0.89 (mean = 0.36, s.d. = 0.15), and the number of shared species ranging from zero to 11 (mean = 3.30, s.d. = 1.83).

Wild bee species richness increased with plant species richness and with the proportion of semi-natural habitat edges in the surrounding landscape (Table 1, Fig. 2). The goodness of fit of the Poisson GLMs with the proportion of all semi-natural habitat edges improved gradually with the size of the buffer within which edge habitat was estimated. The lowest AICc value was obtained from the model with a 1500m buffer (Table 1). Forest edges were an important contributor to the value of seminatural habitat edges as bee habitat. Indeed, considering only forest edges yielded models of bee species richness that were comparable to the models that included all edge habitats (|ΔAICc| = 1.2) but better than the null model that included only plant species richness (|ΔAICc| < 2). By contrast, excluding forest edges reduced the goodness of fit compared to models that included all semi-natural habitat edges (|ΔAICc| = 3) or only forest edges (|ΔAICc| = 4.3).

**Table 1.** Comparisons of Poisson GLMs of wild bee species richness as a function of semi-natural edge habitats within circular buffers around sites.

| Wild bee species richness | AICc<sub>Semi-natural</sub> | AICc<sub>Forest</sub> | AICc<sub>Non-forest</sub> |
| --- | --- | --- | --- |
| Edges in 250m radius + plant richness | 371.1 | 371.7 | 368.1 |
| Edges in 500m radius + plant richness | 372.1 | 371.2 | 371.0 |
| Edges in 750m radius + plant richness | 370.7 | 368.8 | 371.9 |
| Edges in 1km radius + plant richness | 369.2 | 367.6 | 372.0 |
| Edges in 1.25km radius + plant richness | 369.3 | 367.8 | 372.0 |
| Edges in 1.5km radius + plant richness | 368.8 | 367.6 | 371.8 |
| Wild bee species richness<sub>null model</sub> | AICc<sub>Null model</sub> |  |  |
| Plant species richness | 369.9 |  |  |

*\*Comparisons were done using AICc values across models where edge habitat was identified* *using all semi-natural edge habitats (AICc<sub>Semi-natural</sub>), forest edges only (AICc<sub>Forest</sub>), or all* *non-forest edge habitats (AICc<sub>Non-forest</sub>). Poisson GLMs of wild bee species richness were* *compared to a null model that only included plant species richness (AICc<sub>Null model</sub>).*

**Figure 2.**
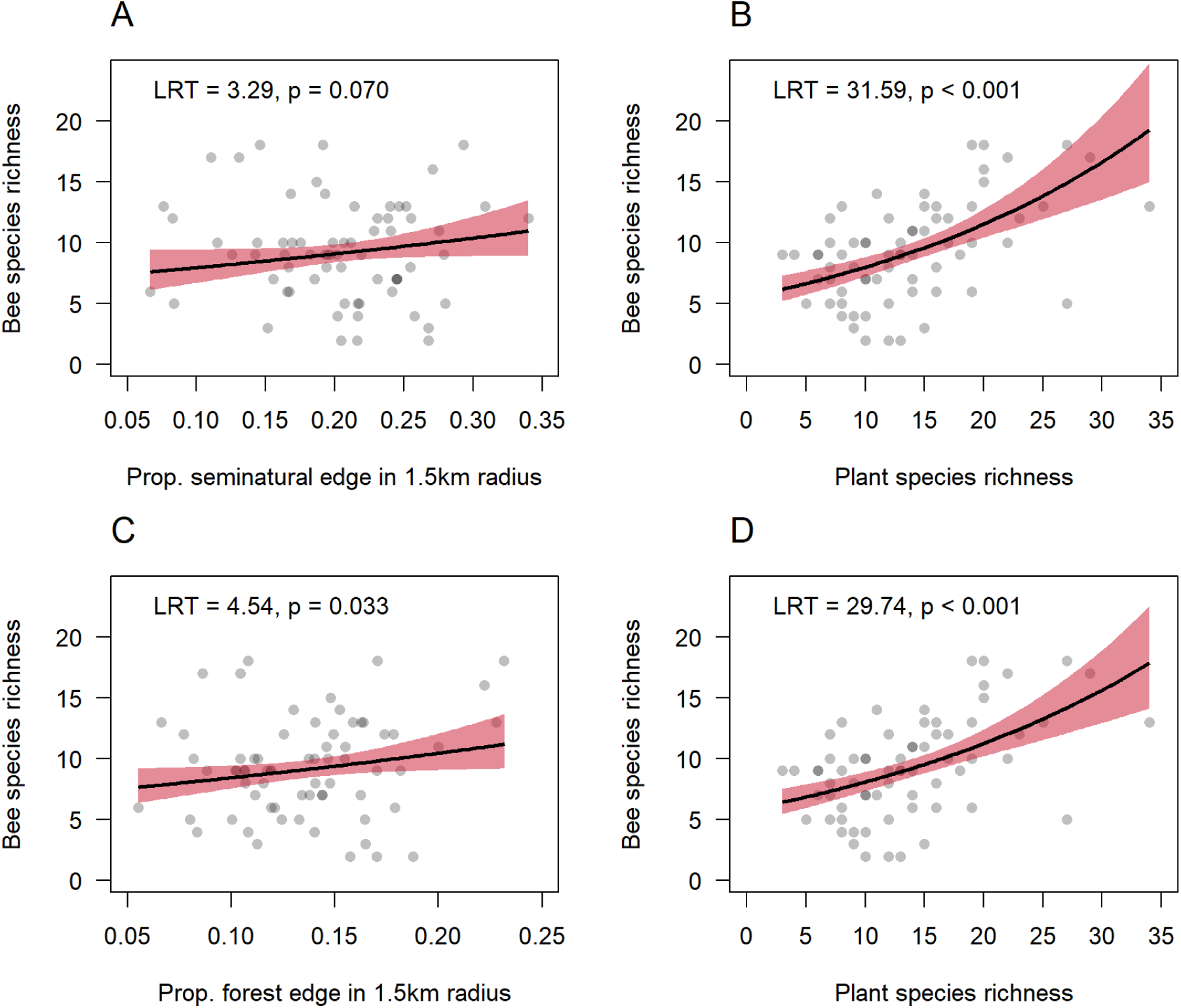
Marginal effects plots showing that wild bee species richness increased with the proportion of semi-natural habitat edges within 1.5km of sampling sites (A), after controlling for the increase in bee species richness with plant species richness (B), and that this relationship was mainly driven by an increase in be species richness with the proportion of forest edges (C) after controlling for local plant species richness (D).

Wild bee species compositional similarity decreased with the length of least cost paths (Table S1) identified from the semi-natural habitat edge map (Fig. 3A, d.f. = 1, χ^2^ = 33.9, p < 0.001) which provided a slightly better fit (|ΔAICc| = 1.3, Table S2) than that of the least cost path lengths when considering only forest edges (d.f. = 1, χ^2^ = 32.6, p < 0.001). Both zero-inflated GLMMs with least cost path lengths provided better models (|ΔAICc| > 3.9, Table S2) of wild bee species compositional similarity than the model that included geographic distance (d.f. = 1, χ^2^ = 28.7, p < 0.001). Contrary to our expectations, models of bee compositional similarity as functions of the least cost path costs did not perform better than the geographic distance model (|ΔAICc| < 2, Table S2). Bee species compositional similarity was highest between site-pairs that both had a high plant species richness and lowest when comparing a plant species rich site to a plant species poor site (Fig. 3B, plant richness site i × plant richness site j interaction term: d.f. = 1, χ^2^ = 17.21, p < 0.001). Results were broadly similar when assessing connectivity as the number of shared species between site-pairs with the least cost path length models outperforming (|ΔAICc| > 2.7, Table S2) and least cost path cost models performing similarly or even slightly worse (|ΔAICc| = 2.1) to the geographic distance model. The decrease in shared species was modelled equally well (|ΔAICc| = 1, Table S2) by the seminatural edge habitat least cost path model (d.f. = 1, χ^2^ = 27.0, p < 0.001) and the forest edge least cost path model (d.f. = 1, χ^2^ = 26.1, p < 0.001). As for species compositional similarity, the number of shared species between site-pairs depended on the plant species richness of the two sites (Fig. 3D, plant richness site i × plant richness site j interaction term: d.f. = 1, χ^2^ = 19.63, p < 0.001). See Table S1 for full zero-inflated Beta GLMM and generalized Poisson GLMM model summary statistics.

**Figure 3.**
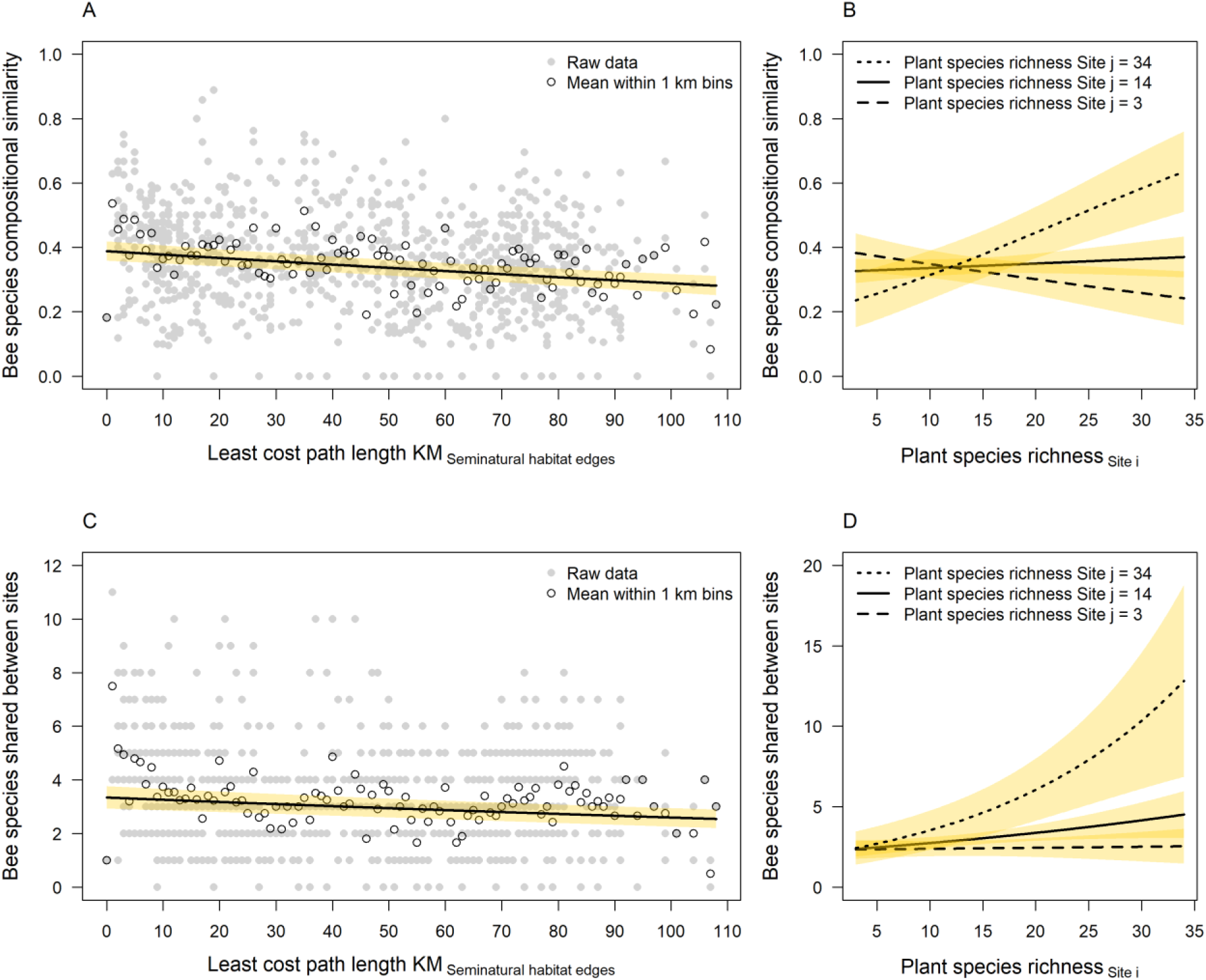
Marginal effects plots showing that bee species compositional similarity between pairs of sites decreased with the length of least cost path dispersal corridors (A) while also depending on the plant species richness within the two sites being compared (B). Results were similar when assessing connectivity as the number of shared bee species between site-pairs (C-D).

We used the map of our study region (Fig. 4A) with the proportion of seminatural grasslands (Fig. 4B) to identify least cost paths between seminatural grasslands of conservation priority in Norway (Fig. 4C) and Denmark (Fig. 4D). Mapping the least cost paths revealed how Norwegian seminatural grasslands that for historical reasons are often located in forested areas can be viewed as a network of habitats where connectivity can be improved by targeting habitat improvement schemes to seminatural edges that connect grassland habitats. For the Danish case the prioritized grassland habitats occurred in clusters, defined by the Natura2000 target areas for 2022-2027. The least cost paths revealed how corridors between clusters of prioritized habitats are likely to converge along shared paths where habitat improvement schemes are likely to improve the connectivity between multiple habitats.

**Figure 4.**
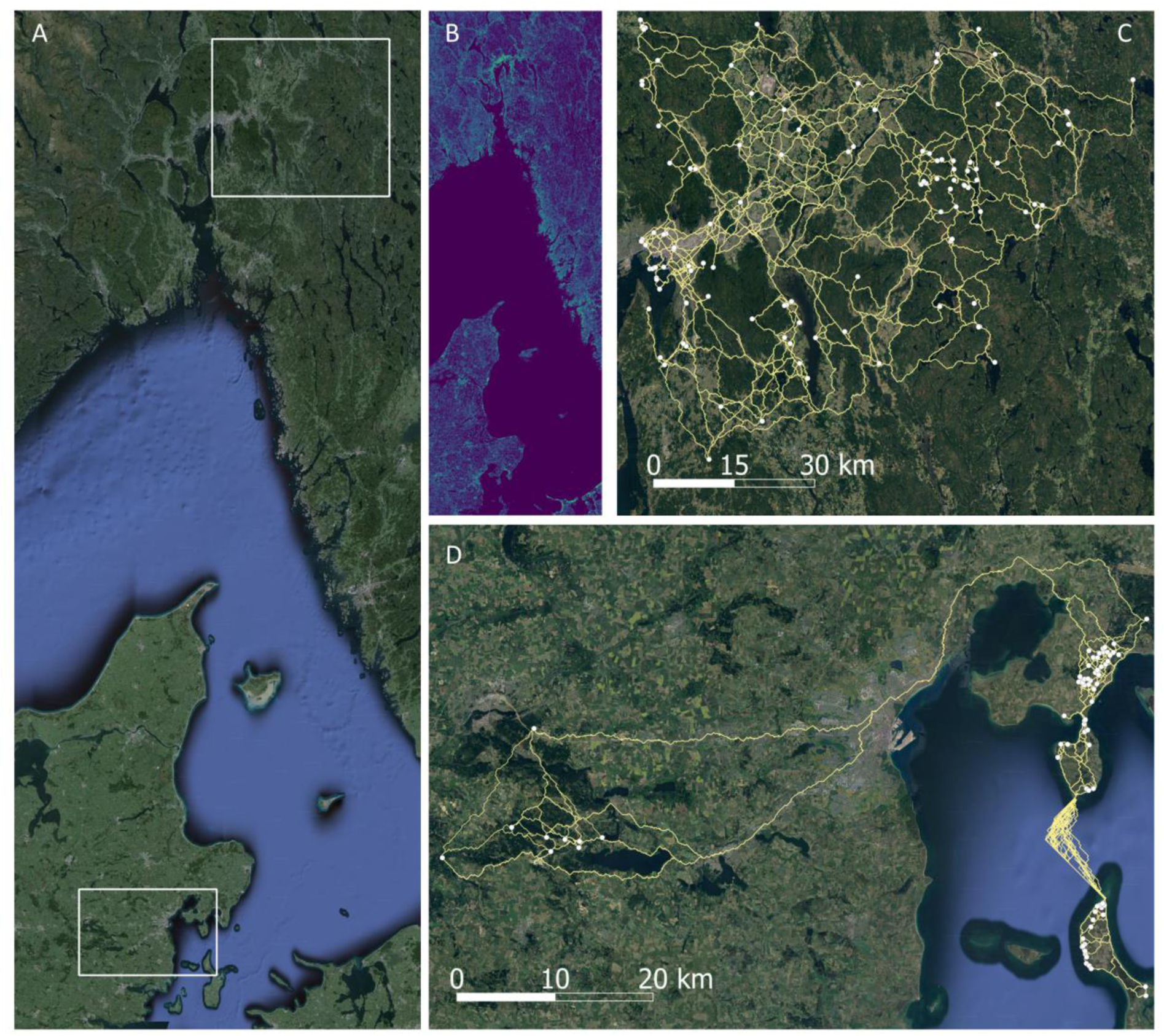
Overview map of the Norwegian study region showing the extents of the two study landscapes (A). Within the study region we used a map showing the proportionate contribution of semi-natural habitat-edges within 100m grid cells (B) to identify least cost path corridors between seminatural grasslands (white points) of conservation priority in the Norwegian (C) and Danish (D) study landscapes. Satellite imagery from Map data ©2024 Google via QGIS 2024.

## Discussion

We found that the presence of seminatural habitat edges not only increased species richness within bee communities but also the connectivity of between bee communities. Least cost path lengths between site-pairs provided models with a better fit to the species compositional similarity and to differences in the number of shared species, than models using inter-site geographical distances suggesting that seminatural habitat edges act as dispersal corridors in the landscape. However, least cost paths were substantially longer than the geographical distances, stressing the importance of habitat quality for bee movement across the landscape.

Seminatural habitat edges may increase dispersal rates by providing wild bees with foraging and nesting habitats along dispersal routes. Habitat resources are often a limiting factor for bee species diversity in intensively managed agricultural (Le Féon et al., 2010; Steffan-Dewenter et al., 2002) and forested (Winfree et al., 2007) landscapes. In forested landscapes, clear-cuts can provide important floral resources for pollinators (Rubene et al., 2015; Nielsen and Totland 2014), however clear-cuts quickly lose their value as habitat due to regrowth and canopy closure (Zitomer et al., 2023). That bee diversity is often lower in forested micro-habitat sites compared to clear-cuts, or forest edges (Mullally et al., 2019), suggests that the edge habitat - and not the forest itself - provides resources for bees. In forested landscapes, seminatural edge habitats occur in the form of sun-exposed forest edges (Sydenham et al. 2014, Proesmans et al. 2019), road verges (Eckerter et al., 2023) and power line clearings (Eldegard et al., 2015, Wagner et al., 2019) which provide important, and persistent, habitat resources for wild bees. Indeed, and in line with our findings, the species richness and abundance of flower-visitors to oilseed rape (Bailey et al. 2014) and coffee (Ricketts 2004) has been shown to decrease with distances to forest edges. However, wild bees require large quantities of pollen to sustain their populations, with many bee species requiring the pollen from more than 30 flowers to rear a single larva (Müller et al., 2006). Because of the high pollen requirements of many species, the short foraging range of wild bees (Zurbuchen et al. 2010), and the limited size of semi-natural habitat edges which restricts the amounts resources they can provide (Johansen et al. 2022), semi-natural habitat edges are unlikely to be able to sustain large bee populations for extended periods of time. Seminatural habitat edges should therefore not be viewed as a substitute for seminatural grassland habitats (Von Königslöw et al. 2021).

While seminatural habitat edges may not be able to replace seminatural grasslands (Von Königslöw et al. 2021), they may act as important stepping-stones (Menz et al. 2011), increasing the dispersal rate between seminatural grasslands that are of conservation concern. In forested landscapes, patch colonization experiments have shown that strips of open, non-forested habitat can act as corridors by facilitating colonization (Griffin & Haddad 2021) as has been also shown in urban settings (Balbi et al. 2019, 2021). However, the functioning of semi-natural habitat edges as dispersal corridors for bees likely depends on the floral and nesting resources they provide. Indeed, we found that the bee species richness within sites increased with local plant richness, indicated by bee communities from plant species rich sites sharingmore species. In contrast, bee communities on plant species poor sites or on sites that differed in their plant diversity shared fewer bee species. The importance of plant species richness in relation to the habitat quality of edges has also been found for bumblebee and butterfly species richness along Swedish roadsides (Horstmann et al. 2023). Using maps of least-cost paths between new, restored, or existing semi-natural grassland habitats can provide a useful tool for identifying and directing management efforts towards potentially important dispersal corridors. Facilitating the movement of wild bees across landscapes through directed management interventions that improve habitat conditions along forest edges is likely to increase the rate at which species disperse and colonize suitable habitats.

Management schemes for improving habitat conditions for wild bees along seminatural habitat edges will differ depending on the type of habitat edge. For roadsides traffic safety restricts the type of management schemes that can be adopted and where they can be implemented. In Norway for instance, vegetation must be repeatedly mowed during the season along curved roads, near road crosses, and near bus stops, for safety reasons to ensure visibility (Pers. Comm. Astrid B. Skrindo). For roadsides along linear road sections management regimes could entail mowing parts of the edge habitat, to prevent them becoming overgrown with grasses, while leaving other parts as refuges (Buri et al. 2014). Allowing for grass tussocks to form along parts of the roadside can improve the nesting conditions for bumblebees where several species nest under thick swards of grass (O’Connor et al. 2017). For the mowed parts, the hay should ideally be removed after mowing (Jakobsson et al., 2018). The timing of mowing should be tailored to the flowering time of the local flora to ensure floral resources are present when bee require them, and that plants are given time to produce and disperse their seeds (Johansen et al., 2019). Altered mowing regimes are however only likely to give immediate effects if plants have already established a seedbank or seedlings. Indeed, the natural establishment of species rich plant communities along roadsides can take several decades (Horstmann et al. 2023). The plant diversity in roadsides without a species rich seedbank can be improved by sowing seeds from pollen and nectar producing plants. Seed-mixes that are often used contain non-native plants which should be avoided because of the potential threat they may pose to the native flora (Valkó et al. 2023). Also, the presence of non-native, and invasive, plants may require the road authorities to intensify mowing regimes (Pers. Comm. Astrid B. Skrindo). An alternative is to use seed-rich hay collected from local seminatural grasslands and spread the hay to allow seeds to establish along roadsides. The ‘hay-method’ has previously been shown to be effective and less costly than using commercial seeds when establishing seminatural grasslands (Rydgren et al. 2010) and comes with the added benefit that it contributes to conserving local plant genetic diversity. In addition to improving plant species richness and bumblebee nesting conditions through altered mowing regimes, nesting sites for ground nesting bees can be introduced in the form of non-vegetated sand pockets (Fortel et al. 2016). Improving pollinator habitat through altered mowing regimes, potentially coupled with the hay-method, and the establishment of nesting sites for ground nesting bees can also be adopted along other edge habitats, such as grassland and forest edges. Along these other types of habitat-edges it is also possible to cater for cavity nesting bees by allowing dead wood to gather (Westerfelt et al. 2015), and by preserving or building stone drywalls (Xie et al., 2020). Implementing these management actions will incur a cost for managers and should be directed to where they will have the desired effect. Maps of least cost paths between pollinator habitats of conservation priority can be used to identify potentially important dispersal corridors along which management actions should be prioritized.

We used the compositional similarity and numbers of shared species between sites as measures of connectivity. The compositional similarity between sites is known to depend on the sample size within site-pairs, and under sampling may inflate estimates of (dis-)similarity (Beck et al. 2013, Stier et al. 2016). Still, despite that our sampling effort was restricted to three surveys per site, and that of the variability in local plant species richness which was strongly related to bee species richness and compositional similarity, we were still able to detect a statistically significant relationship between least cost path lengths and habitat connectivity between sites. These findings support those of Sydenham et al., (2017) that the species diversity within bee communities can be dispersal limited. That habitat conditions in the landscape can increase the dispersal rate of bees has been shown in agricultural landscapes, where high amounts of seminatural habitat reduces the distance decay in species compositional similarity (Beduschi et al. 2018). However, to our knowledge it has not previously been shown that least cost paths lengths are more strongly related to bee community connectivity than geographic distance.

Our results suggest that linear edge habitats contribute to the connectivity among bee communities in intensively managed temperate landscapes. However, the value of such edges as habitat depends on their plant diversity. While implementing management actions such as altered mowing regimes to increase plant diversity seminatural habitat edges is costly least-cost-path analyses can provide an efficient tool for identifying important dispersal corridors along which management actions should be prioritized. Considering dispersal corridors as an integrated part of pollinator conservation planning will allow the establishment of networks of high-quality habitats where the longevity of populations of pollinators can be maintained through dispersal dynamics.

## Acknowledgements

We are grateful to Gry Liljefors, Daniel J. Skoog, Jens Mogens Olesen, Kaj-Andreas Hanevik, Lise Lauridsen, and Anders Gunnar Helle, for assistance with fieldwork and to Arnstein Staverløkk for identifying bumblebees sampled in Norway. Viken county roads administration and John W. Dirksen from Eidskog municipality contributed to identifying appropriate study sites in Norway.

## Author contributions

**Markus A. K. Sydenham**: Conceptualization (lead); Data curation (lead); Formal analysis (lead); Funding acquisition (lead); Investigation (Lead); Methodology (lead); Project administration (lead); Validation (lead); Visualization (lead); Writing – original draft (lead); Writing – review and editing (lead). **Anders Nielsen:** Conceptualization (supporting); Funding acquisition (supporting); Writing – review and editing (supporting). **Yoko L. Dupont:** Data curation (co-lead); Funding acquisition (supporting); Investigation (co-lead); Methodology (supporting); Writing – review and editing (supporting). **Henning B. Madsen:** Investigation (supporting); Data curation (supporting); Writing – review and editing (supporting). **Claus Rasmussen:** Funding acquisition (supporting); Methodology (supporting); Writing – review and editing (supporting). **Marianne S. Torvanger**; Investigation (supporting); Data curation (supporting); Writing – review and editing (supporting). **Bastiaan Star:** Conceptualization (supporting); Writing – review and editing (supporting)

## Data archiving statement

Data and R code associated with this paper will be d}eposited on Dryad if the manuscript is accepted for publication.

## Ethics statement

Alle fieldwork was conducted following national regulations.

## Supplementary material S1

**Table S1.**
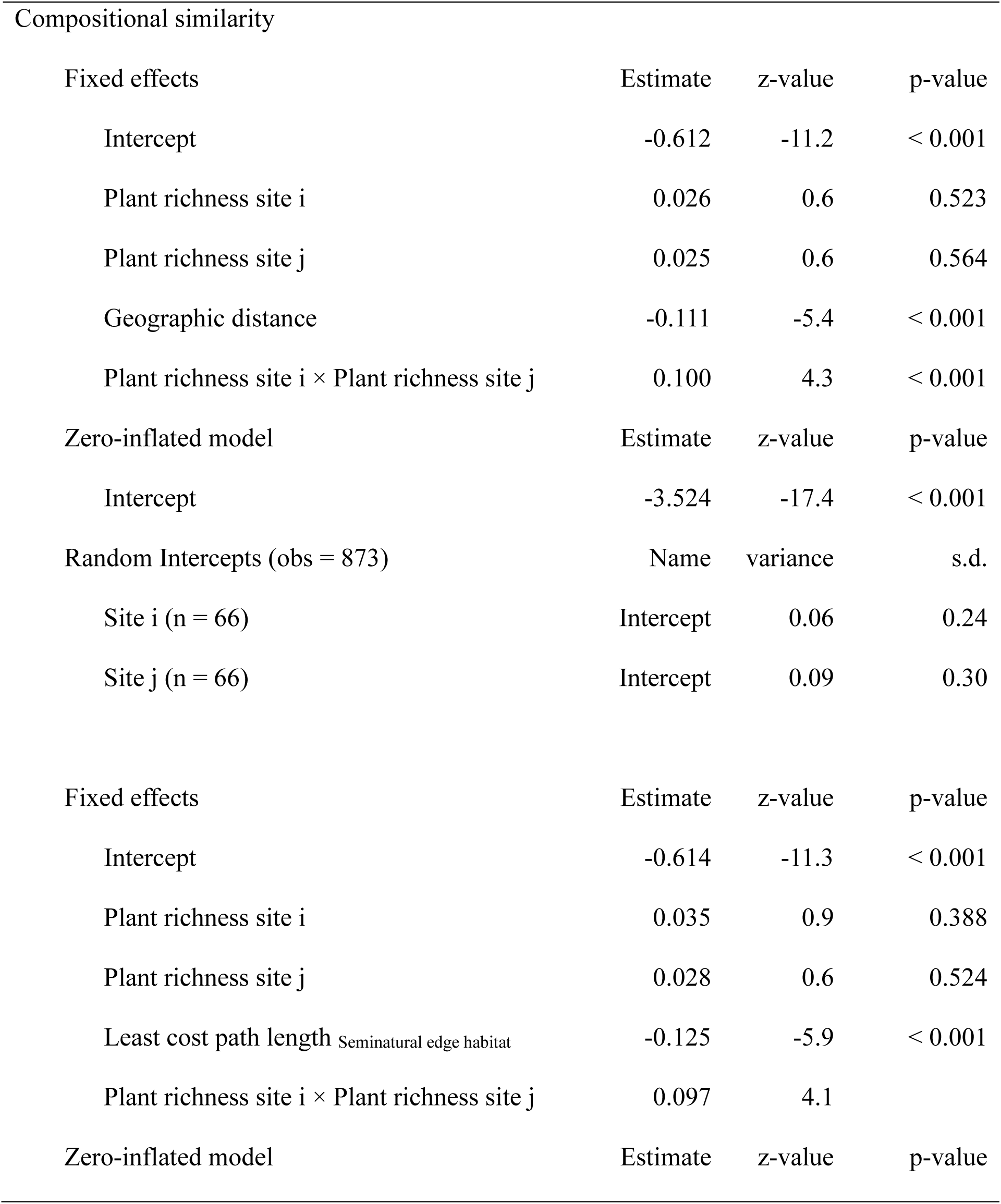
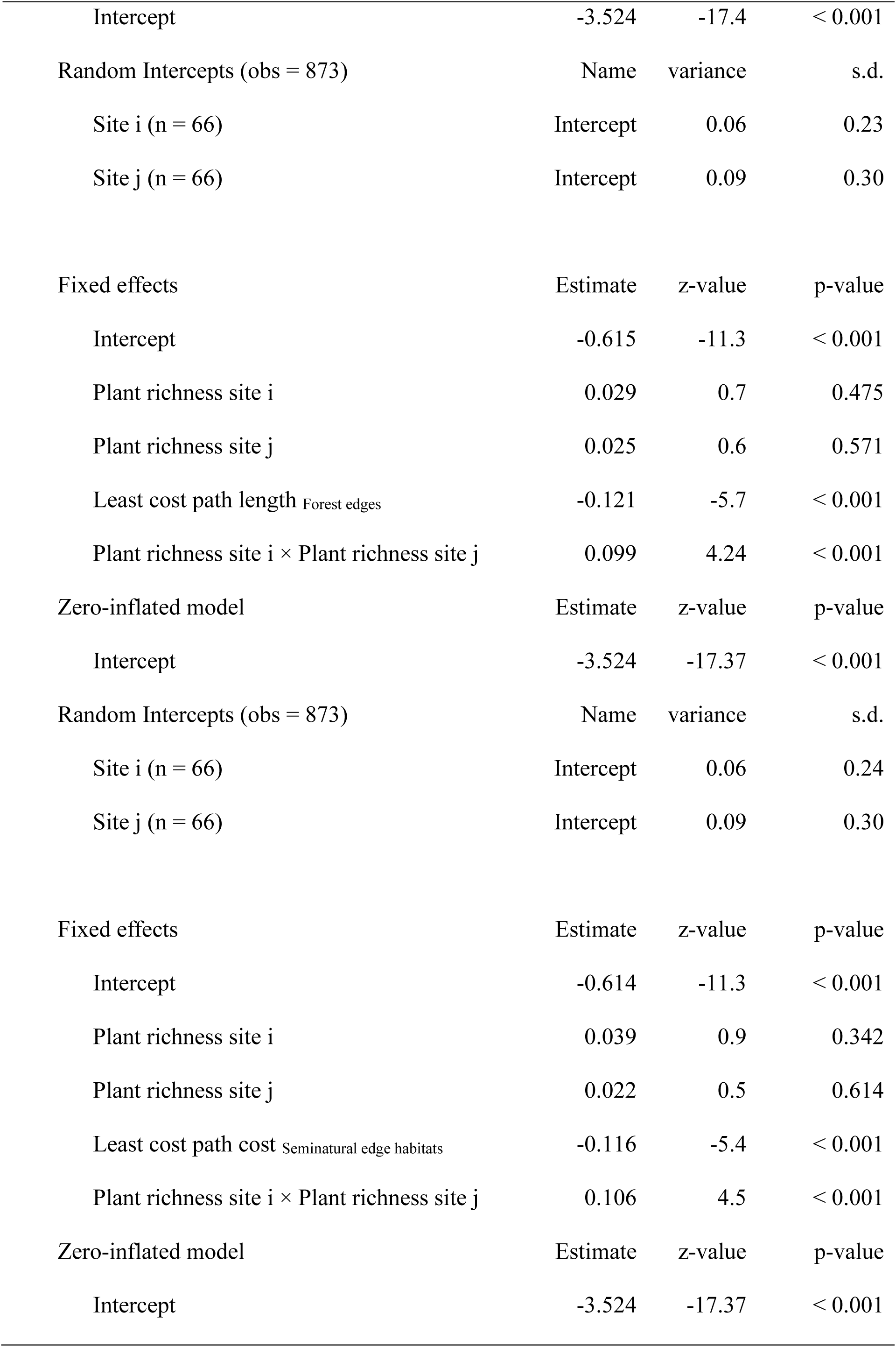
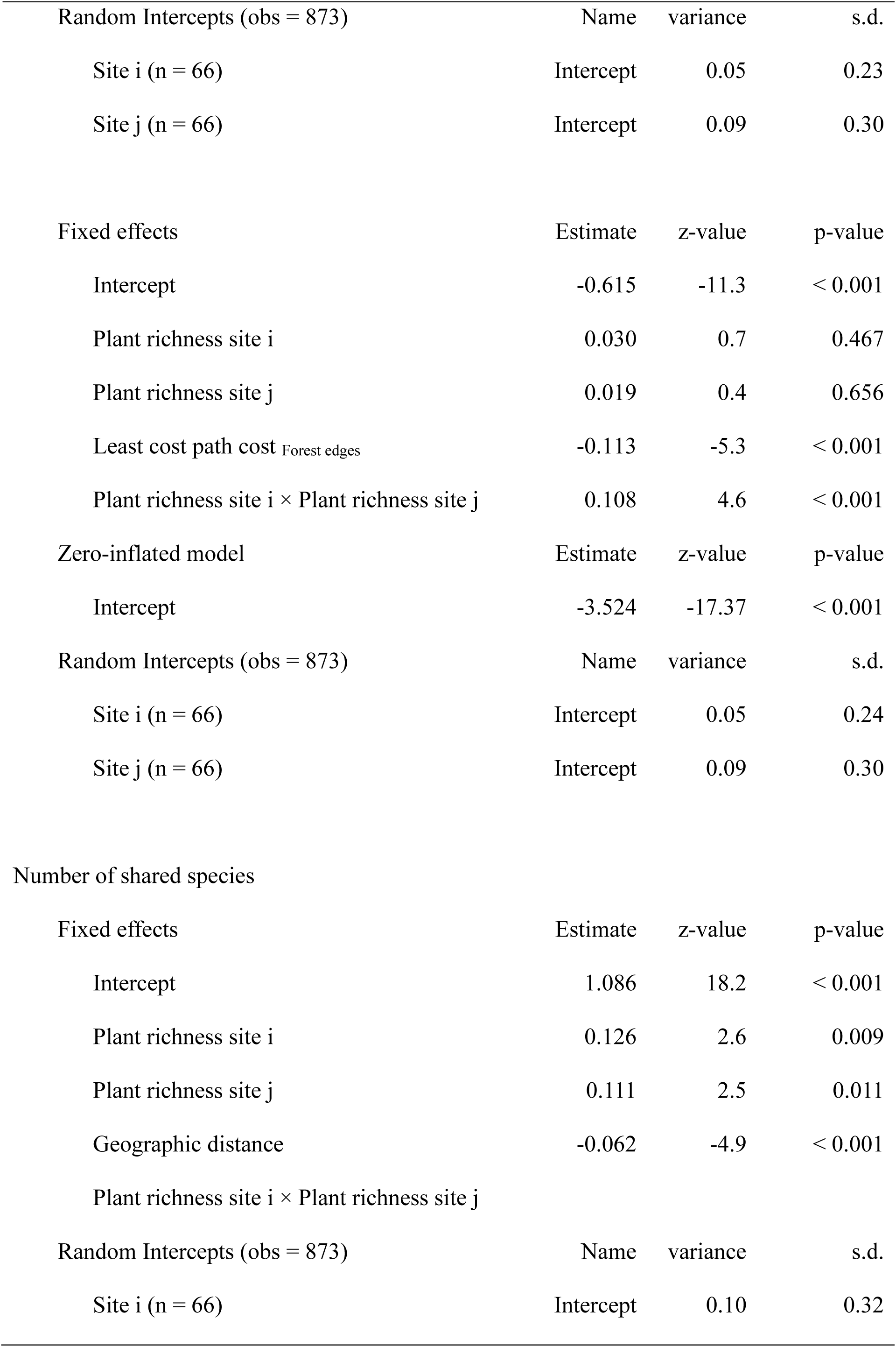
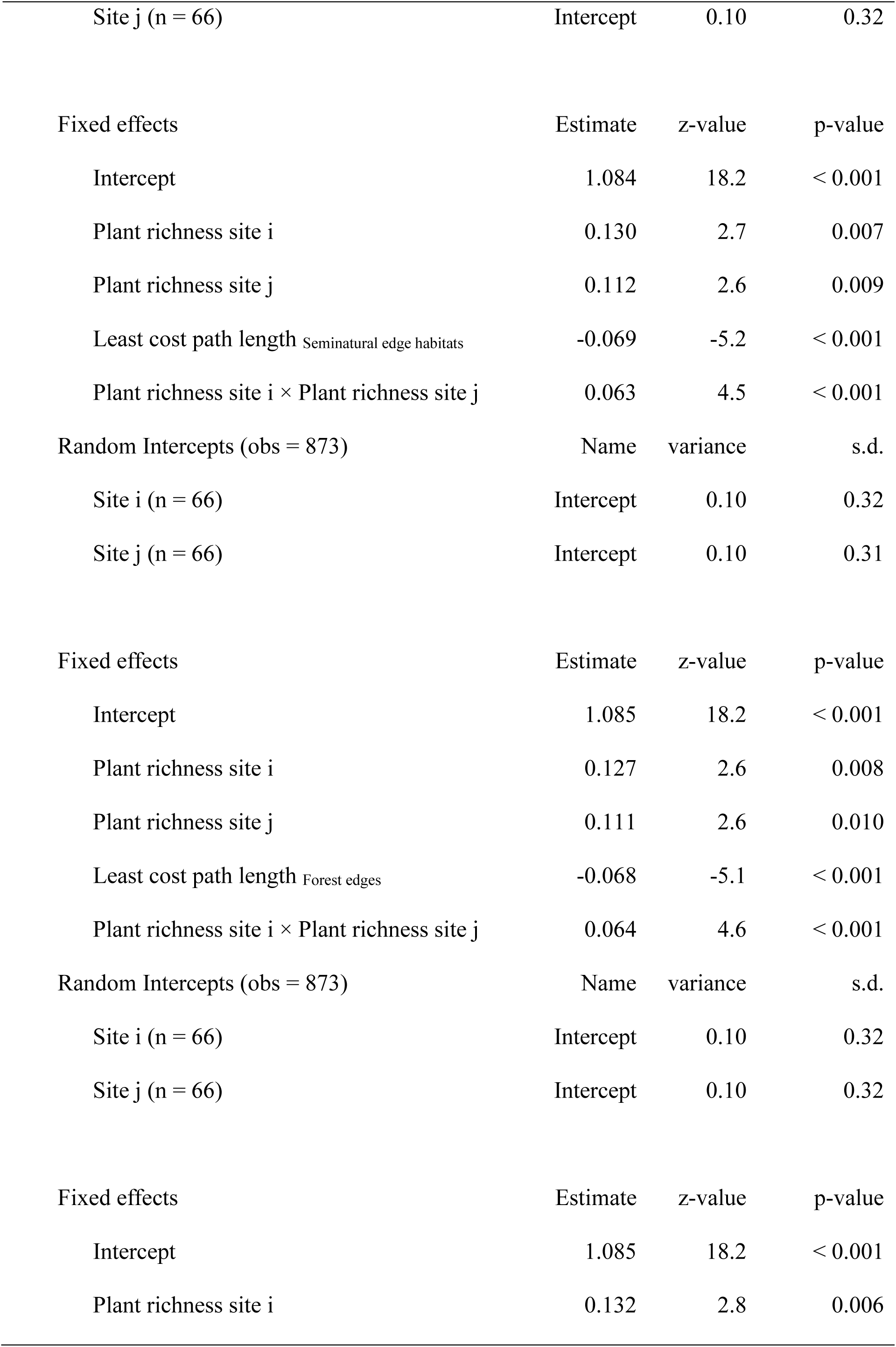
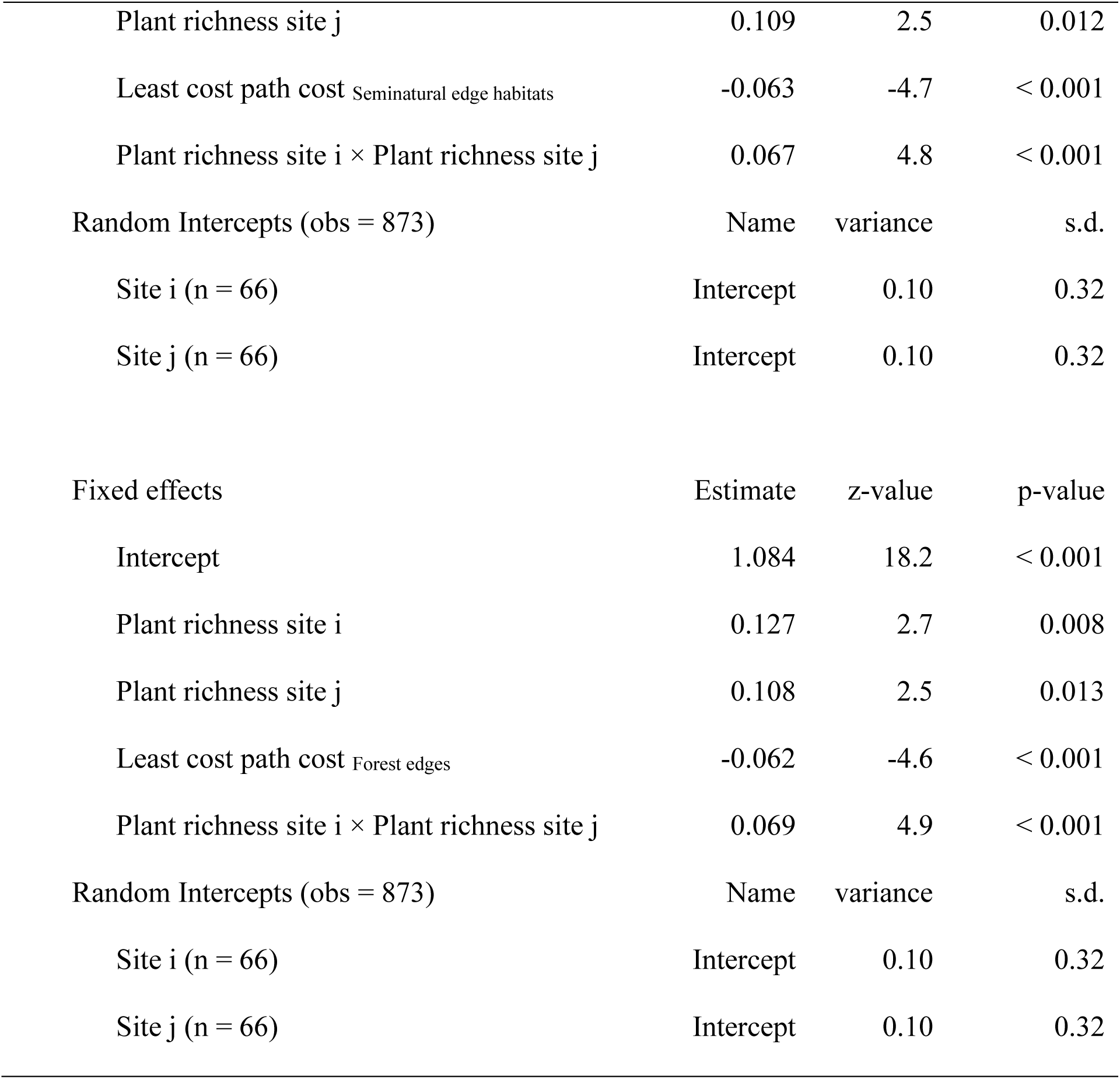
Summary outputs from Zero-inflated Beta GLMMs used to model wild bee species compositional similarities between sites and from Generalized Poisson GLMMs used to model the number of shared species between sites.

| Compositional similarity |  |  |  |  |
| --- | --- | --- | --- | --- |
| Fixed effects |  | Estimate | z-value | p-value |
| Intercept |  | -0.612 | -11.2 | < 0.001 |
| Plant richness site i |  | 0.026 | 0.6 | 0.523 |
| Plant richness site j |  | 0.025 | 0.6 | 0.564 |
| Geographic distance |  | -0.111 | -5.4 | < 0.001 |
| Plant richness site i × Plant richness site j | | 0.100 | 4.3 | < 0.001 |
| Zero-inflated model |  | Estimate | z-value | p-value |
| Intercept |  | -3.524 | -17.4 | < 0.001 |
| Random Intercepts (obs = 873) |  | Name | variance | s.d. |
| Site i (n = 66) |  | Intercept | 0.06 | 0.24 |
| Site j (n = 66) |  | Intercept | 0.09 | 0.30 |
| Fixed effects |  | Estimate | z-value | p-value |
| Intercept |  | -0.614 | -11.3 | < 0.001 |
| Plant richness site i |  | 0.035 | 0.9 | 0.388 |
| Plant richness site j |  | 0.028 | 0.6 | 0.524 |
| Least cost path length Seminatural edge habitat |  | -0.125 | -5.9 | < 0.001 |
| Plant richness site i × Plant richness site j | | 0.097 | 4.1 | |
| Zero-inflated model |  | Estimate | z-value | p-value |

|  |  |  |  |
| --- | --- | --- | --- |
| Intercept | -3.524 | -17.4 | < 0.001 |
| Random Intercepts (obs = 873) | Name | variance | s.d. |
| Site i (n = 66) | Intercept | 0.06 | 0.23 |
| Site j (n = 66) | Intercept | 0.09 | 0.30 |
| Fixed effects | Estimate | z-value | p-value |
| Intercept | -0.615 | -11.3 | < 0.001 |
| Plant richness site i | 0.029 | 0.7 | 0.475 |
| Plant richness site j | 0.025 | 0.6 | 0.571 |
| Least cost path length Forest edges | -0.121 | -5.7 | < 0.001 |
| Plant richness site i × Plant richness site j | 0.099 | 4.24 | < 0.001 |
| Zero-inflated model | Estimate | z-value | p-value |
| Intercept | -3.524 | -17.37 | < 0.001 |
| Random Intercepts (obs = 873) | Name | variance | s.d. |
| Site i (n = 66) | Intercept | 0.06 | 0.24 |
| Site j (n = 66) | Intercept | 0.09 | 0.30 |
| Fixed effects | Estimate | z-value | p-value |
| Intercept | -0.614 | -11.3 | < 0.001 |
| Plant richness site i | 0.039 | 0.9 | 0.342 |
| Plant richness site j | 0.022 | 0.5 | 0.614 |
| Least cost path cost Seminatural edge habitats | -0.116 | -5.4 | < 0.001 |
| Plant richness site i × Plant richness site j | 0.106 | 4.5 | < 0.001 |
| Zero-inflated model | Estimate | z-value | p-value |
| Intercept | -3.524 | -17.37 | < 0.001 |
| Random Intercepts (obs = 873) | Name | variance | s.d. |
| Site i (n = 66) | Intercept | 0.05 | 0.23 |
| Site j (n = 66) | Intercept | 0.09 | 0.30 |
| Fixed effects | Estimate | z-value | p-value |
| Intercept | -0.615 | -11.3 | < 0.001 |
| Plant richness site i | 0.030 | 0.7 | 0.467 |
| Plant richness site j | 0.019 | 0.4 | 0.656 |
| Least cost path cost Forest edges | -0.113 | -5.3 | < 0.001 |
| Plant richness site i × Plant richness site j | 0.108 | 4.6 | < 0.001 |
| Zero-inflated model | Estimate | z-value | p-value |
| Intercept | -3.524 | -17.37 | < 0.001 |
| Random Intercepts (obs = 873) | Name | variance | s.d. |
| Site i (n = 66) | Intercept | 0.05 | 0.24 |
| Site j (n = 66) | Intercept | 0.09 | 0.30 |
| Number of shared species |  |  |  |
| Fixed effects | Estimate | z-value | p-value |
| Intercept | 1.086 | 18.2 | < 0.001 |
| Plant richness site i | 0.126 | 2.6 | 0.009 |
| Plant richness site j | 0.111 | 2.5 | 0.011 |
| Geographic distance | -0.062 | -4.9 | < 0.001 |
| Plant richness site i × Plant richness site j | | | |
| Random Intercepts (obs = 873) | Name | variance | s.d. |
| Site i (n = 66) | Intercept | 0.10 | 0.32 |
| Site j (n = 66) | Intercept | 0.10 | 0.32 |
| Fixed effects | Estimate | z-value | p-value |
| Intercept | 1.084 | 18.2 | < 0.001 |
| Plant richness site i | 0.130 | 2.7 | 0.007 |
| Plant richness site j | 0.112 | 2.6 | 0.009 |
| Least cost path length Seminatural edge habitats | -0.069 | -5.2 | < 0.001 |
| Plant richness site i × Plant richness site j | 0.063 | 4.5 | < 0.001 |
| Random Intercepts (obs = 873) | Name | variance | s.d. |
| Site i (n = 66) | Intercept | 0.10 | 0.32 |
| Site j (n = 66) | Intercept | 0.10 | 0.31 |
| Fixed effects | Estimate | z-value | p-value |
| Intercept | 1.085 | 18.2 | < 0.001 |
| Plant richness site i | 0.127 | 2.6 | 0.008 |
| Plant richness site j | 0.111 | 2.6 | 0.010 |
| Least cost path length Forest edges | -0.068 | -5.1 | < 0.001 |
| Plant richness site i × Plant richness site j | 0.064 | 4.6 | < 0.001 |
| Random Intercepts (obs = 873) | Name | variance | s.d. |
| Site i (n = 66) | Intercept | 0.10 | 0.32 |
| Site j (n = 66) | Intercept | 0.10 | 0.32 |
| Fixed effects | Estimate | z-value | p-value |
| Intercept | 1.085 | 18.2 | < 0.001 |
| Plant richness site i | 0.132 | 2.8 | 0.006 |
| Plant richness site j | 0.109 | 2.5 | 0.012 |
| Least cost path cost Seminatural edge habitats | -0.063 | -4.7 | < 0.001 |
| Plant richness site i × Plant richness site j | 0.067 | 4.8 | < 0.001 |
| Random Intercepts (obs = 873) | Name | variance | s.d. |
| Site i (n = 66) | Intercept | 0.10 | 0.32 |
| Site j (n = 66) | Intercept | 0.10 | 0.32 |
| Fixed effects | Estimate | z-value | p-value |
| Intercept | 1.084 | 18.2 | < 0.001 |
| Plant richness site i | 0.127 | 2.7 | 0.008 |
| Plant richness site j | 0.108 | 2.5 | 0.013 |
| Least cost path cost Forest edges | -0.062 | -4.6 | < 0.001 |
| Plant richness site i × Plant richness site j | 0.069 | 4.9 | < 0.001 |
| Random Intercepts (obs = 873) | Name | variance | s.d. |
| Site i (n = 66) | Intercept | 0.10 | 0.32 |
| Site j (n = 66) | Intercept | 0.10 | 0.32 |

**Table S2.**
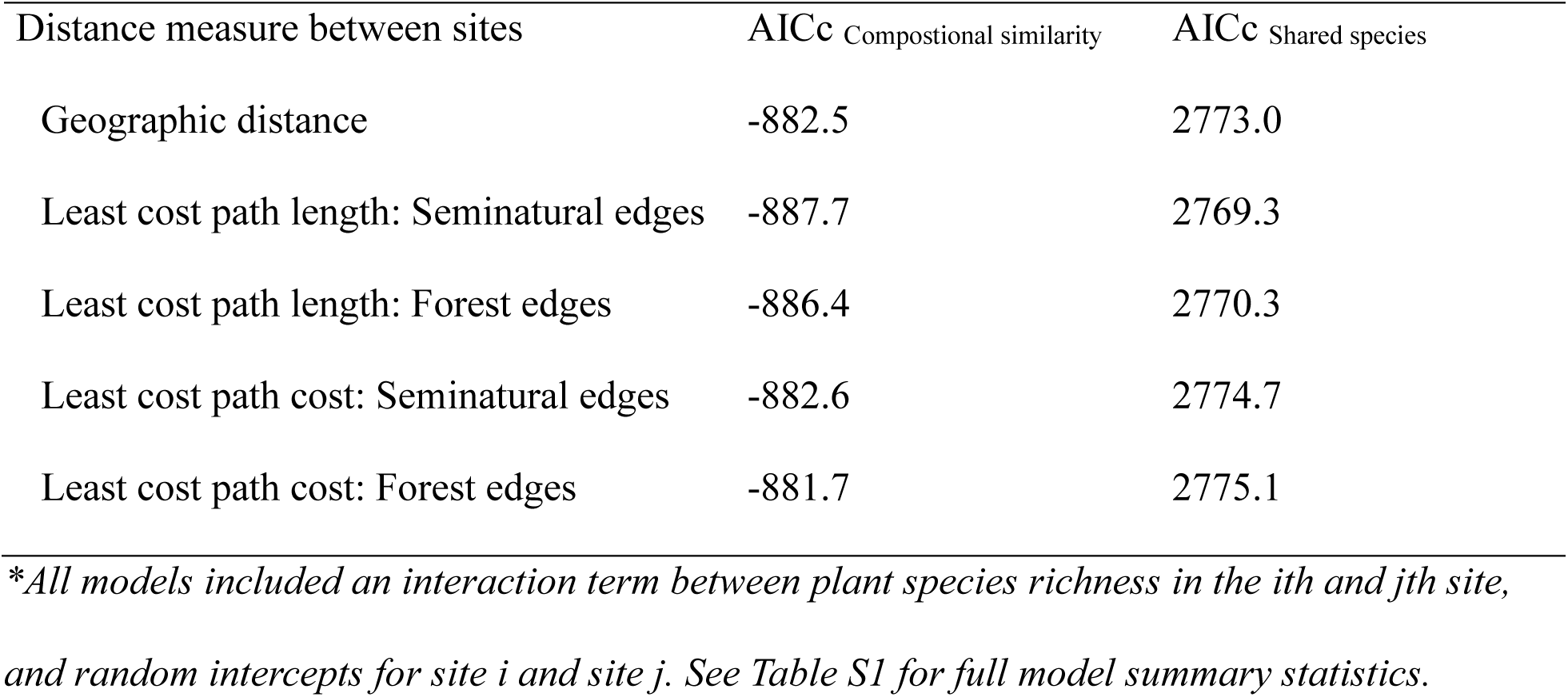
Comparisons of zero-inflated Beta GLMMs of wild bee species compositional similarity or generalized Poisson GLMMs of the number of shared species richness between site-pairs *i* and *j* as a function of geographic distance, seminatural edge habitat least cost path length, forest edge least cost path length, seminatural edge habitat least cost path cost, and forest edge least cost path cost.

| Distance measure between sites | AICc Compositional similarity | AICc Shared species |
| --- | --- | --- |
| Geographic distance | -882.5 | 2773.0 |
| Least cost path length: Seminatural edges | -887.7 | 2769.3 |
| Least cost path length: Forest edges | -886.4 | 2770.3 |
| Least cost path cost: Seminatural edges | -882.6 | 2774.7 |
| Least cost path cost: Forest edges | -881.7 | 2775.1 |

*\*All models included an interaction term between plant species richness in the $i$ th and $j$ th site,* *and random intercepts for site $i$ and site $j$ . See Table S1 for full model summary statistics.*

**Figure S1.**
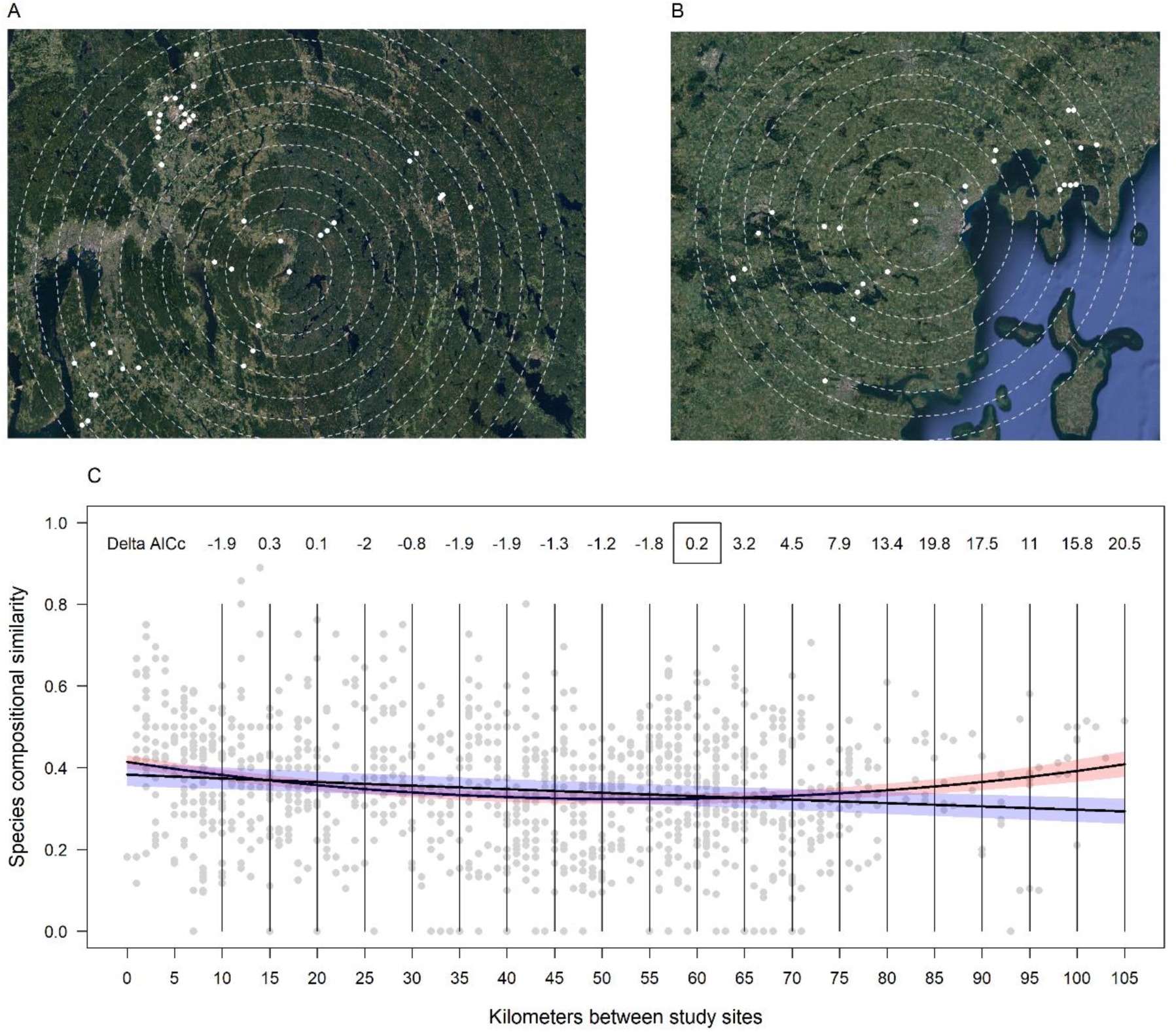
Location of study sites in Norway (A) and Denmark (B) shown within buffers of increasing 5km distance from the most central site within each region. Zero-inflated Beta GLMMs with km between sites untransformed or using a second order polynomial were used to model bee species compositional similarity between sites and compared using AICc values (C). For analyses restricted to site-pairs within 60km of each other the untransformed and the polynomial models performed equally well (ΔAICc = 0.2), while at greater distances the polynomial model performed best (ΔAICc > 2). Species compositional similarity was calculated as 1 – the bray-curtis dissimilarity. Regression lines in (C) show fitted relationships ± 2×standard deviations for the untransformed (blue polygon) and polynomial models (red polygon) including all site distances.

